# Iso-propyl stilbene: A life-cycle signal?

**DOI:** 10.1101/498857

**Authors:** Alexia Hapeshi, Jonatan Mimon Benarroch, David James Clarke, Nicholas Robin Waterfield

**Affiliations:** Microbiology and Infection Unit, Division of Biomedical Sciences, Warwick Medical School, University of Warwick; School of Microbiology and APC Microbiome Institute, University College Cork, Cork, Ireland

**Keywords:** 3,5-dihydroxy-4-isopropyl-*trans*-stilbene, *Photorhabdus*, secondary metabolites

## Abstract

Members of the Gram-negative bacterial genus *Photorhabdus* are all highly insect pathogenic and exist in an obligate symbiosis with the entomopathogenic nematode worm *Heterorhabditis*. All members of the genus produce the small molecule 3,5-dihydroxy-4-isopropyl-*trans*-stilbene (IPS) as part of their secondary metabolism. IPS is a multi-potent compound that has antimicrobial, antifungal and anticancer activities and also plays an important role in symbiosis with the nematode. In this study we have examined the response of *Photorhabdus* itself to exogenous ectopic addition of IPS at physiologically relevant concentrations. We observed that the bacteria had a measureable phenotypic response, which included a decrease in bioluminescence and pigment production. This was reflected in changes in its transcriptomic response, in which we reveal a reduction in transcript levels of genes relating to many fundamental cellular processes, such as translation and oxidative phosphorylation. Our observations suggest that IPS plays an important role in the biology of *Photorhabdus* bacteria, fulfilling roles in quorum sensing, antibiotic-competition advantage and maintenance of the symbiotic developmental cycle.

## INTRODUCTION

Stilbenoids are natural products with important biological properties. Most are synthesised by plants in response to pathogenic attack. However, *Photorhabdus* and *Bacillus* bacteria engaged in symbioses with entomopathogenic nematodes also produce certain stilbene derivatives (1,2). *Photorhabdus* is an entomopathogenic bacterium, which forms a symbiotic relationship with *Heterorhabditis* nematodes. Infection of insect larvae by the nematodes results in regurgitation of the bacteria into the insect (3). *Photorhabdus* then produces a large set of toxins and digestive enzymes to kill and utilise the insect as a food source (4). Additionally, it releases multiple antimicrobial factors to eliminate competition in the insect carcass by other bacteria or fungi found in the soil (5,6). 3,5-dihydroxy-4-isopropyl-*trans*-stilbene (IPS) (figure 1) is a small molecule produced by *Photorhabdus*, which has been shown to have antimicrobial (1), antifungal (7,8) and anticancer (9) activities and also seems to play an important role in symbiosis (10).

**Figure 1.**
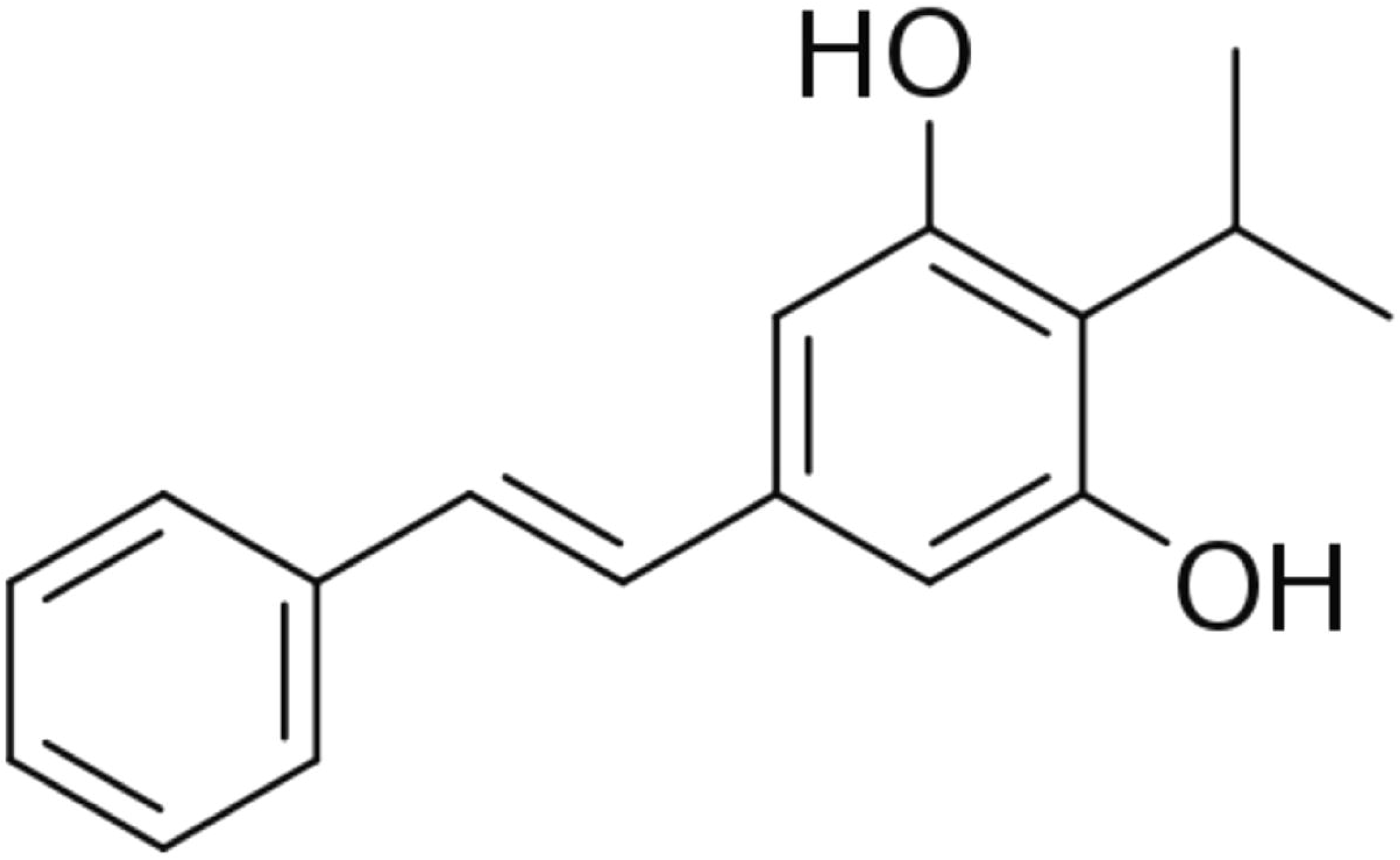
The structure of 3,5-dihydroxy-4-isopropyl-*trans*-stilbene (IPS)

IPS is produced by or encoded in the genomes of all *Photorhabdus* strains investigated so far (1,7,8,11–13). Additionally, depending on the strain and the growth phase of the bacteria other stilbene derivative compounds may also be synthesised (1,8). Joyce et al (2008) demonstrated that *Photorhabdus* utilises a pathway distinct from that used in plants for IPS biosynthesis. The *Photorhabdus* biosynthetic pathway is composed of two branches that converge, with one originating in phenylalanine metabolism and the other in branched-chain amino acid metabolism. Additionally, one of the stilbene precursors is involved in the production of branched chain fatty acids and it is thus possible that fatty acid oxidation can contribute to IPS production.

In vitro, the expression of *stlA*, which encodes a phenylalanine-ammonium lyase that catalyses the first step in stilbene biosynthesis is induced by nutrient limitation (14). Activity of the *stlA* promoter seems to negatively correlate with the amount of nutrients in the media and shows distinct changes at the different growth phases (14,15). The production of the alarmone molecules (p)ppGpp is required for this as deletion of both *relA* and *spoT* results in a shutdown of *stlA* promoter activity (16). Additionally, post-transcriptional regulation plays a role in controlling *stlA* expression and this is mediated via the BarA/UvrY two-component system and CsrA/*csrB* (17). However, a *uvrY* deficient strain showed no significant difference in the actual amounts of stilbenes produced (18). Moreover, it has been established that the production of IPS is repressed by the LysR-type regulator HexA that controls the synthesis of secondary metabolites in general (18,19) and is itself regulated by the action of Hfq (20). This illustrates that multiple levels of regulation are present which affect the levels of stilbenes.

In vivo, IPS production is associated with entry of the bacteria into the stationary phase of growth following replication in the insect host. In an infection model using *Galleria mellonella* larvae, the production of IPS by *Photorhabdus* is detectable 24 h post infection after which the levels of IPS remain stable for several days (21). It should be noted that by 24 h, the population of *Photorhabdus* within *Galleria* increases to a very high density of 10^9^ bacteria/g of wet larvae (21). Actual concentrations of IPS vary depending on the entomopathogenic complex used and subsequently the *Photorhabdus* symbiont strain. Levels can range from ∼600 to ∼4000 µg/g wet insect tissue (21,22). Additionally, Eleftherianos et al. (2006) have estimated that in the haemolymph of *Manduca sexta* caterpillars that have been infected with *Photorhabdus luminescens* subsp. *laumondii* TT01 the levels of IPS can reach up to 275 – 550 µg/ml (15).

One suggested function of the stilbenes, as a result of their antimicrobial and antifungal properties, is the protection of the insect carcass from competition. Another possible role is in the establishment of a successful symbiosis. A *Photorhabdus* stilbene-negative mutant is severely impaired in its ability to promote infective juvenile recovery into adult hermaphrodites in an in vitro symbiosis assay (10). However, the presence of IPS in itself is not sufficient for the induction of IJ recovery (10) indicating that other factors produced by the bacteria are also necessary for the process. With this in mind, we examined the effect of IPS on *P*. *luminescens* itself. It has been previously shown that high concentrations of IPS (>100 µg/ml) are toxic to the bacteria (23) but the mechanism behind this has not been investigated in detail. We demonstrate that the presence of IPS actually has a profound effect on central metabolism, secondary metabolism and consequently the phenotype of the bacteria. We propose that the timing of IPS production in an insect infection plays a crucial role in orchestrating the relationship between the bacteria and the entomopathogenic nematode host.

## RESULTS

In order to investigate the effect of IPS on *Photorhabdus* bacteria we first provided exogenous IPS at various concentrations from the beginning of growth in batch culture. At this growth phase the bacteria do not normally synthesize the compound. Under these conditions *P*. *luminescens* shows a clear dose dependent response to ectopic IPS, with growth rate reducing as the concentration of the compound added increases (figure 2A). A clear dose response in bioluminescence was also observed, the levels of which decreased with higher IPS concentrations. This holds true until the later stages of growth when the levels of bioluminescence, in the presence of IPS, actually become higher than in the control cultures (figure 2B).

**Figure 2.**
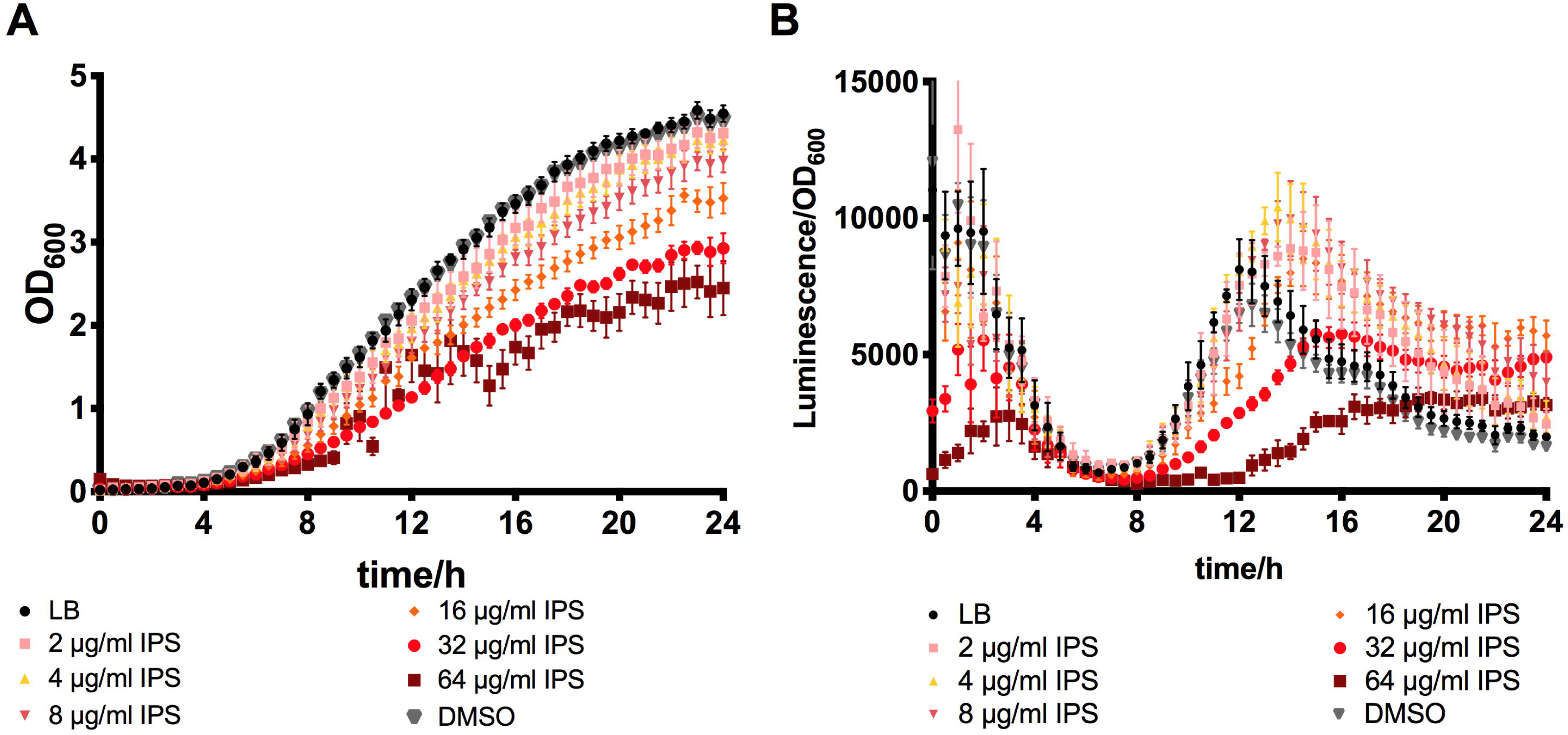
Dose dependent effect of IPS on A) growth and B) bioluminescence of *P*. *luminescens* TT01. Bacteria were grown in LB supplemented with either DMSO or different concentrations of IPS. Shown are the averages of 4 biological replicates from 3 independent experiments ± standard error.

Based on these findings we subsequently chose a concentration of 32 µg/ml and investigated whether addition of IPS to bacteria that were already growing exponentially, for approximately 5 h, would have the same effect as above. Indeed, we observed that early on after treatment there were significantly lower (q value < 0.01) levels of bioluminescence in the cultures that were treated with the compound compared to those treated with the carrier solvent DMSO only. Interestingly, again by 24 h this effect was reversed (q value < 0.001). Furthermore, subsequent growth was also moderately negatively affected, which was significant even at 30 min post treatment (q value < 0.001), with IPS treated cultures having a mean OD_600_ at 600 nm of 0.49, while DMSO treated cultures had a mean OD_600_ of 0.54 (figure 3).

**Figure 3.**
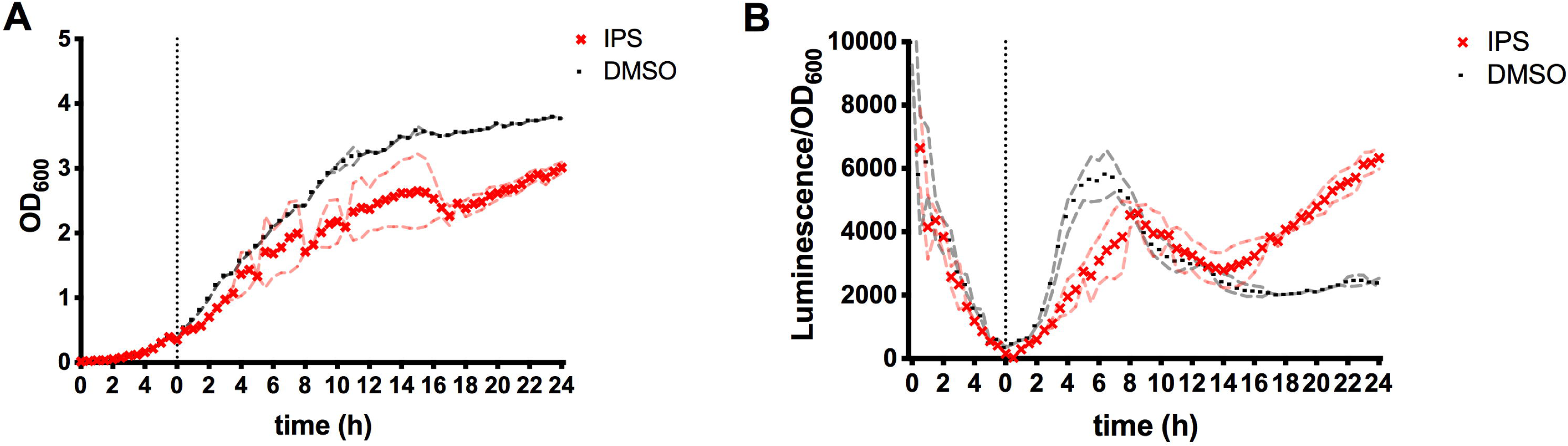
The effect of IPS addition at mid-exponential on growth (A) and luminescence (B) of *P*. *luminescens* TT01. The bacteria were grown for ∼5.5 h and 32 μg/ml IPS was added to the cultures. Shown are the averages from three biological replicates ± standard error.

Additionally, we noted that by the end of the experiment (24h growth), the cultures of *P*. *luminescens* were seen to have very little pigmentation when prematurely exposed to IPS, compared to untreated cultures which showed the typical orange/red pigmentation (figure 4a). This observation was corroborated by spectrophotometric measurements of the cultures (data not shown). *P*. *luminescens* normally produces increasing amounts of anthraquinone pigments at late stages of its growth phase. The absence of the red colour suggests that production of anthraquinones is either directly or indirectly inhibited by IPS. Interestingly in a mutant strain where IPS production is prevented, through disruption of *stlA* by transposon insertion (24), the resulting colonies are hyper-pigmented, also suggesting a deregulation of anthraquinone production (figure 4b).

**Figure 4.**
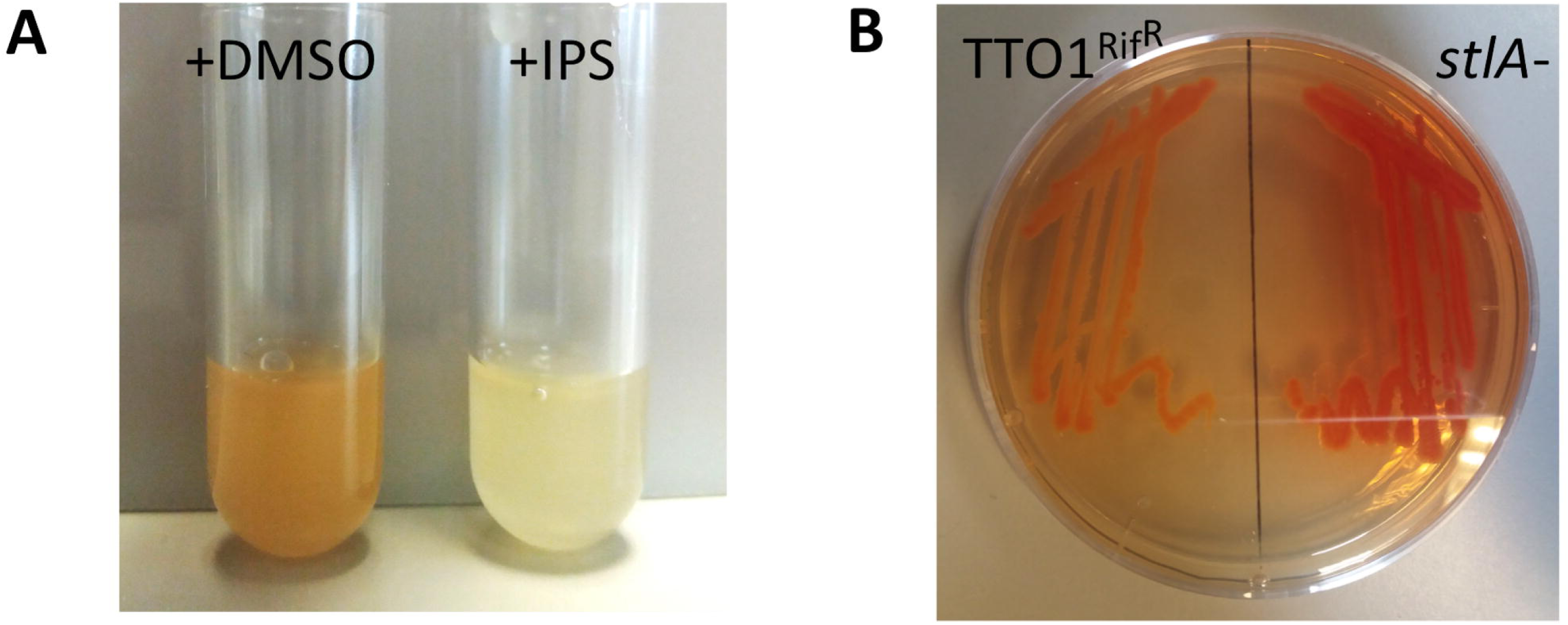
The influence of IPS on pigment production. A) Cultures of *P*. *luminescens* following addition of either the DMSO solvent alone or 32 μg/ml IPS (in DMSO) and growth for 24 h. B) *P*. *luminescens* TTO1 Rif^R^ and a *stlA*^*-*^ mutant strains streaked out on LB agar and incubated for 96 h.

Since IPS appears to affect aspects of *P*. *luminescens* secondary metabolism we decided to investigate its role on the formation of a biofilm by *P*. *luminescens*. The ability to form a biofilm has been implicated in the successful establishment of symbiosis with the *Heterorhabditid* nematode (25) and may be part of a pathogenic / symbiotic life-cycle switch in *Photorhabdus*. Furthermore, previous work has associated *Photorhabdus* secondary metabolites with a role in nematode symbiosis (20,26). We performed this experiment using LB media as this was previously shown to support better biofilm formation in *P*. *temperata* than BHI or SOC media (27). It should be noted that wild-type *Photorhabdus* does not form very thick biofilms in vitro, and the biofilms they do form in standard microplate assays appear to occur mostly on the liquid-air interface. Supplementation of the cultures with IPS did not have a significant effect on the total amount of biofilm formed (figure 5a). However, when this assay was repeated with the *stlA* mutant, which forms a much thicker biofilm, addition of IPS resulted in a significant reduction in the amount of biofilm suggesting that IPS plays a modulatory role (figure 5b).

**Figure 5.**
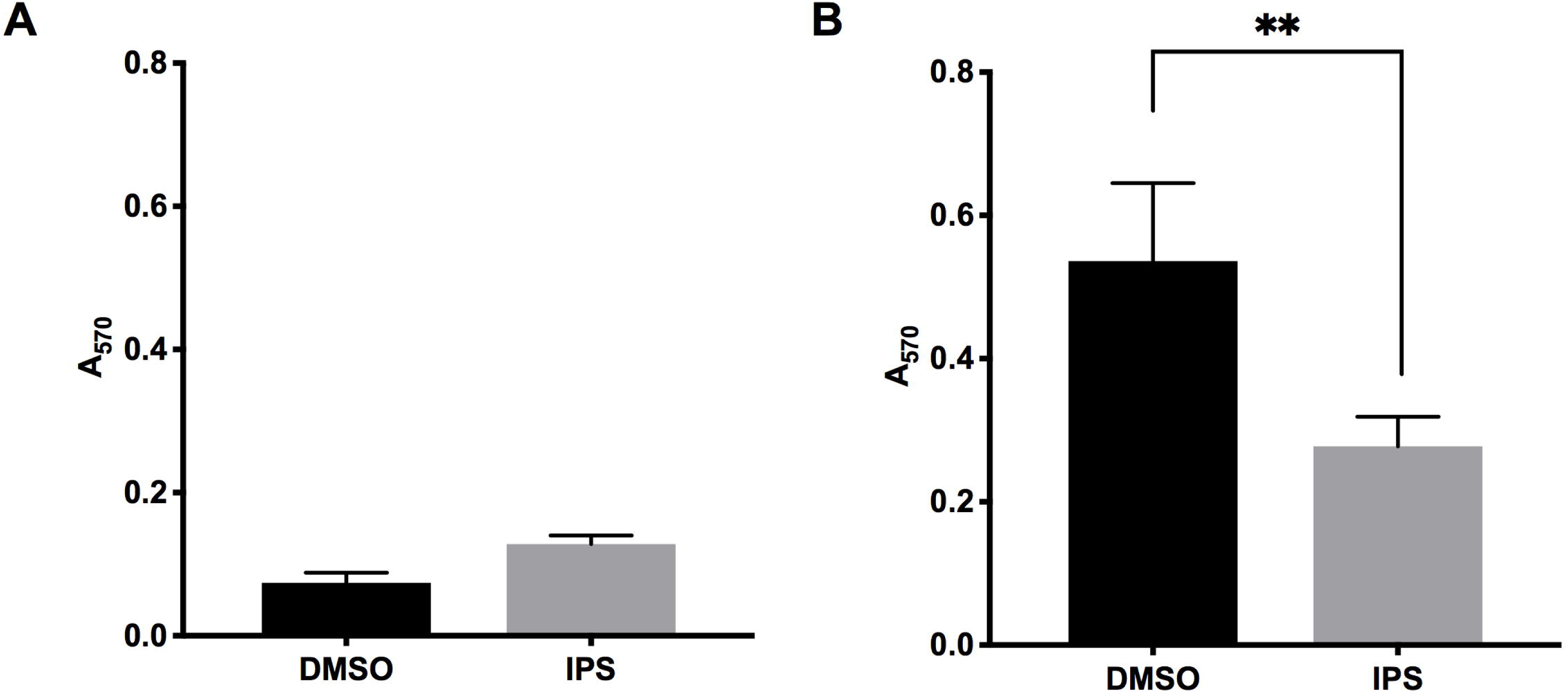
The effect of IPS on *P*. *luminescens* biofilm. The amount of biofilm formed by A) *P*. *luminescens* TTO1 Rif^R^ and B) *P*. *luminescens stlA-* as measured using a crystal violet assay in the presence of either the DMSO solvent or 32 μg/ml IPS in DMSO. Results are the average means of four biological replicates ± standard error. ** *p* < 0.01.

In order to elucidate whether the phenotypic observations followed a more systemic pattern we performed an RNA-seq transcriptomic study on *P*. *luminescens* TT01 following addition of exogenous IPS to mid log planktonic cells. At 32 µg/ml of IPS we were able to see a detectable growth delay and reduction in bioluminescence. We therefore used this concentration to investigate the cause of the growth delay, which we reasoned would be reflected in changes in gene expression. Furthermore, *P*. *luminescens* TT01 would be expected to be exposed to at least this IPS concentration (and likely higher) during insect infection (see above). For this experiment IPS was again added to cultures at exponential phase (OD_600_ ∼ 0.45) and samples for RNA extraction were harvested 30 min later. Bioluminescence measurements confirmed the inhibitory effect of the IPS in the samples used for RNA extraction (Figure S1).

The response of *Photorhabdus* to IPS was systemic with 258 genes significantly up-regulated and 463 genes down-regulated, when only considering the p_adj_ threshold (table S1). Of those, 52 genes had at least 2-fold higher expression whilst 84 genes had at least 2-fold lower expression in the cultures treated with IPS compared to those treated with DMSO. For the purposes of this analysis, we also used the small RNA identification tool toRNAdo (28), to identify putative non-coding RNAs and examine if their expression changes in the presence of IPS. From the output of toRNAdo, 80 sRNAs were added to the annotation of the genome and included in the differential gene expression analysis. Of those, 4 were significantly overexpressed and 10 had significantly lower levels of expression in the presence of IPS. One putative sRNA was among the top 20 overexpressed genes (log_2_ fold change of 2.0). This is located opposite a tRNA-Phe, whose levels however appear to be unchanged. Interestingly, we observed that 21 tRNA encoding-sequences were moderately down-regulated following treatment and so were various tRNA modifying enzymes. This is intriguing as it was recently shown that tRNAs can get rapidly degraded shortly after starvation (29) or stress (30), presumably to prevent mistranslation and the accumulation of damaged or misfolded proteins. No tRNAs were up-regulated in the presence of IPS.

### Secondary metabolism

Consistent with our previous observations we observed IPS dependent down-regulation of the transcription of certain genes involved in secondary metabolism. In particular, the biosynthetic gene cluster *antA-antI* (*plu4186-4194*) encoding the type II polyketide synthase and modifying enzymes involved in anthraquinone production (31) were the most down-regulated genes in our dataset. This explains the absence of a pigment in IPS treated cultures of *Photorhabdus* that was observed (see above). Additionally, among the genes with the most marked reduction in transcript levels were genes belonging to the *Photorhabdus* variable region *plu1434–plu1448* (32), some of which are responsible for antibiotic production. Similarly, *cpmA* and *cpmD*, encoding enzymes involved in carbapenem biosynthesis, were down-regulated. It is of note, however, that contrary to other secondary metabolites that are most highly produced at stationary phase, expression of the *cpm* genes is actually maximal during exponential phase (5). Another highly down-regulated cluster of genes is that located in the region *plu1409-plu1415*. Perhaps surprisingly, the expression of the IPS biosynthesis genes *stlA, stlB, stlCDE* and *bkdABC* was not significantly reduced. Note, these genes are encoded across the genome rather than in a single operon, suggesting the pathway regulon was not affected. However, as this was during a time point when the bacteria would be producing low amounts of IPS, if any, it is possible that these genes cannot be repressed further or that there is no feedback acting to control their expression. On the other hand, transcript levels for *plu3042* and *plu3043*, which encode a transaminase and a prephenate dehydratase respectively, are significantly reduced. The products of these genes are likely to be involved in the production of 2,5-dihydrophenylananine, which is the precursor to another cryptic stilbene derivative, 2,5-dihydrostilbene (33). Contrary to our phenotypic observations that addition of IPS causes a rapid drop in bioluminescence the transcript levels of the *lux* operon itself were not affected. Previous work has also suggested that bioluminescence activity appears to be under post-transcriptional control (19). Also, in contrast to the processes discussed above, which were repressed by IPS, transcription of some other secondary metabolite genes was actually up-regulated. For example, this was the case with *plu3533*, encoding a non-ribosomal peptide synthase (NRPS) that potentially produces a gramicidin/tyrocidin antibiotic.

### Basic metabolic processes

In large part the effect of ectopic IPS addition on the bacterial transcriptome resembles a classic stringent response (Figure 6). In many bacteria this is reflected in the up-regulation of amino acid biosynthesis, with the concurrent down-regulation of many aspects of ribosome biogenesis and protein translation (34,35). In particular, in our dataset 52 out of 55 genes encoding proteins that form the ribosomal subunits, or are otherwise associated with ribosomes, were significantly down-regulated. RNA polymerase subunits also exhibit reduced transcript levels. Furthermore, a large number of genes encoding enzymes involved in the Tri-Carboxylic Acid (TCA) cycle are also negatively affected. The response to IPS seen in the transcriptome is however more extensive than we would expect from stringent response alone. Specifically we also see repression of the transcription of genes involved in Oxidative Phosphorylation, including subunits of the F-type ATPase, succinate dehydrogenase and NADH dehydrogenase. In addition, the pathways of purine and pyrimidine metabolism are also strongly affected. In particular, various genes encoding purine and pyrimidine *de novo* biosynthetic enzymes are down-regulated by ectopic IPS exposure. On the contrary, the gene that codes for CpdB, which is involved in nucleotide catabolism is up-regulated and so are *plu4417* encoding uridine phosphorylase, *nudE* encoding an ADP compounds hydrolase, and *plu0661* encoding a putative pyrimidine/purine nucleotide 5’-monophosphate nucleosidase.

**Figure 6.**
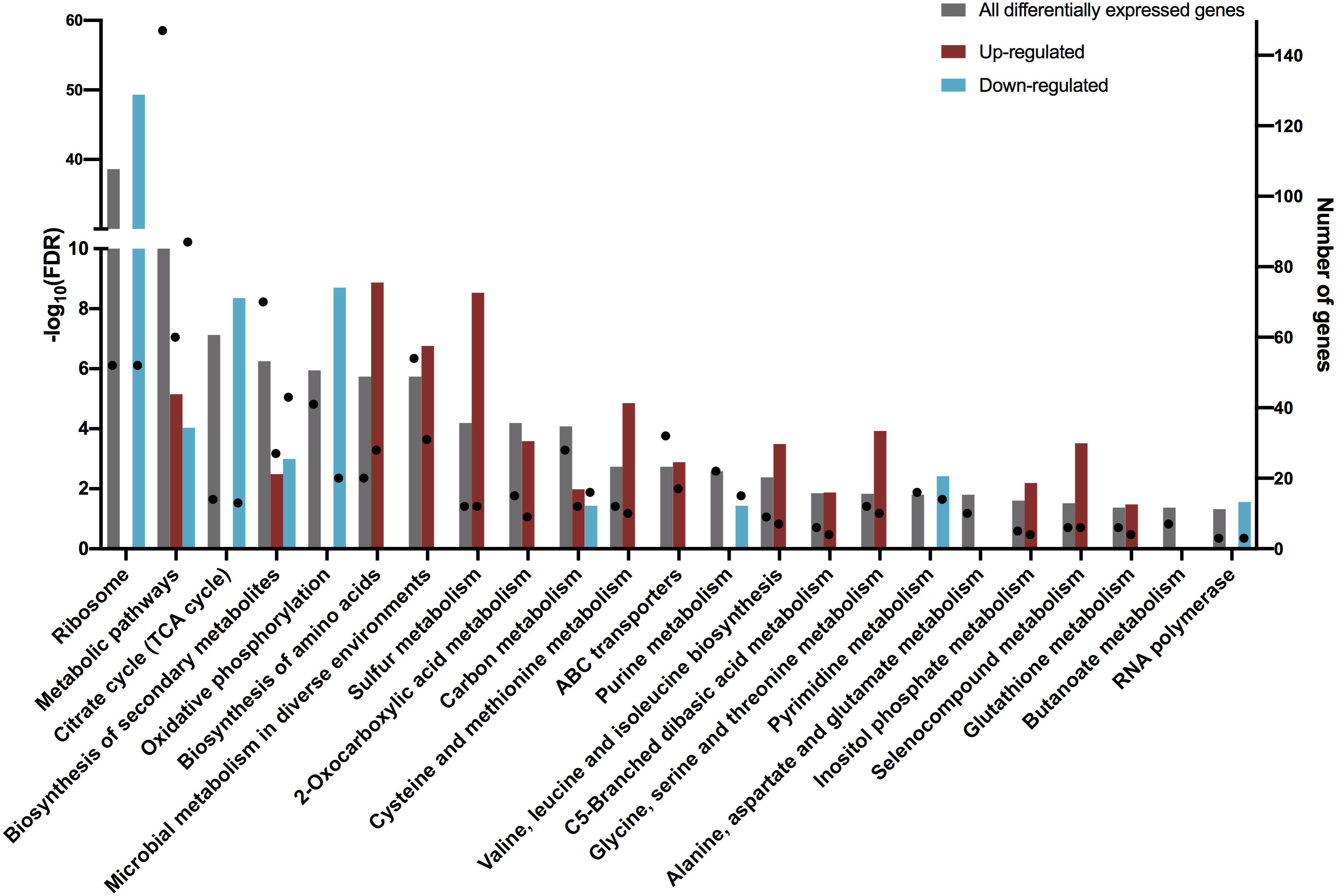
KEGG pathways affected by the presence of ectopic IPS. Overrepresented pathways were identified using STRING version 10.5 using either all the differentially expressed genes in our dataset or the up and down-regulated genes separately. Bars correspond to the –log_10_(FDR), whereby multiple testing correction is done using the Benjamini and Hochberg method (73), and circles indicate the number of affected genes in each pathway.

In other Gram-negative bacteria, the stringent response is controlled by the levels of the (p)ppGpp alarmone, which is in turn regulated by the opposing enzymic activities of the *relA* and *spoT* gene products. Interestingly, we see no change in the transcription of the orthologous *relA* (*plu0910*) and *spoT* (*plu0272*) genes, although this does not rule out any post-transcriptional or post-translational regulation of their activities. To this effect, we observed that the transcript level of *plu4550*, which encodes the essential protein ObgE/CgtA are lower in the presence of IPS. In *E*. *coli* and *Vibrio cholera* CgtA was shown to bind SpoT and it has been suggested that it promotes its (p)ppGpp hydrolase activity (36,37). Hydrolysis of (p)ppGpp would otherwise suppress the stringent response.

Interestingly, amongst the most highly IPS induced genes were those encoding for enzymes involved in sulfur metabolism as well as components of the sulfate/thiosulfate ABC transporter. As the control cultures were treated with an equivalent amount of the IPS carrier solvent DMSO, this cannot be attributed solely to the presence of DMSO in the media. Note however that some of these enzymes are involved in the cysteine and methionine metabolism, which relates to the stringent response-like changes discussed above. Nonetheless, among the overexpressed genes involved in sulfur metabolism was the *sufABCDSE* operon that encodes the Suf pathway proteins, unrelated to amino acid synthesis. The Suf pathway is an iron-sulfur (Fe-S) cluster biogenesis system that operates under stress conditions. SufS is a cysteine desulfurase, which then serves as a donor for the assembly of the Fe-S cluster, which is eventually transferred to other proteins. The Isc system is another Fe-S cluster assembly system and we observed that genes *iscRSUA* encoding the enzymes of the system and the transcription factor IscR, as well as *hscR*, encoding a co-chaperone are also overexpressed. We note that disruption of the TCA cycle in an *mdh* mutant (26) also resulted in an up-regulation of these pathways (unpublished data).

### Transporters

Of note is the significant reduction in transcript levels for *putP*, which encodes a proline (sodium) transporter. On the other hand there was no effect on the expression of *proVXW* encoding the ProU ABC transporter. L-proline plays an important role in the biology of *Photorhabdus*, influencing production of secondary metabolites and affecting the proton motive force. Interestingly, a Δ*putP* mutant had abolished stilbene production but up-regulated anthraquinone (38).

The transcript levels of genes encoding transporters for iron and zinc are increased. For example, *plu3940* and *plu3941* encoding ExbD and ExbB respectively are both overexpressed. Furthermore, genes *plu2320-2321* involved in the biosynthesis of a siderophore related to the yersiniabactin from *Yersinia enterocolitica* are also up-regulated in addition to gene *plu2316*, which encodes a receptor protein that is also predicted to be involved in iron uptake. Iron uptake was previously shown to be important in the virulence of *Photorhabdus*, whilst depending on the bacteria-nematode complex it may also play a role in symbiosis (39,40). On the contrary, the transport of dipeptides, oligopeptides, glutamate/aspartate, arginine and long chain fatty acids is repressed. Similarly, *tctA* and *tctC* that encode subunits of the tripartite tricarboxylate transporter are repressed. We also note that the AcrAB components of the enterobacterial AcrAB-TolC efflux pump are transcriptionally up-regulated. The AcrAB-TolC tripartite system is known to provide resistance to antibiotics but also has a role in pathogenicity (reviewed in (41)). Other affected genes include *tatA* and *tatB* of the twin-arginine translocation system and which are up-regulated, while *secY, secF* and *yidC* that encode components of the Sec pathway of protein export are repressed. Finally, genes *plu0634* and *plu1333* (*rtxB*) encoding subunits belonging to the RTX toxin family of ABC transporters are also repressed.

### Bacterial surface components

Genes *plu2656-plu2660* which belong to the *pgbPE* operon that is required for O-antigen synthesis and assembly (42) are also down-regulated. The *pgbPE* operon is required for *Photorhabdus* virulence as well as colonization of the nematode gut (42). Additionally, a group of genes belonging to the *wbl* cluster whose products are likely to be involved in lipopolysaccharide biosynthesis are down-regulated. These are *wblT, wblR, wblP* and *wblU*. Similarly, genes *plu4658*-*plu4663* encoding enzymes RffG, WecABC and WzzE involved in the synthesis of the enterobacterial common antigen (ECA) are also repressed. This could be significant as in addition to the *pgbPE* operon other genes involved in LPS biosynthesis were also previously shown to be necessary for *Photorhabdus* virulence and transmission to the nematode (43). Furthermore, motility and/or attachment to surfaces may also be affected by the presence of stilbene as we observed a mild drop in transcript levels of *fliT* (*plu1951*), *fliC* (*plu1954*), *fliD* (*plu1953*), encoding flagellar proteins. Additionally, we noted a reduction in transcript levels of *plu00928* and of the operon *plu2156-plu2159*, which are predicted to be involved in the production of fimbriae.

### Transcriptional regulators

A large cluster of genes, which encode PAS4-LuxR solo transcriptional regulators are almost all significantly down-regulated (*plu2001-plu2016*). These are LuxR receptors that lack a cognate LuxI synthase and it has been suggested that they are involved in host hormone-like signal detection (44). Additionally, *plu4444* encoding one of the putative alternative sigma factors FecI, was up-regulated, whilst none of the other homologues of known sigma factors was differentially expressed. This was also the case for *lrp* and *tyrR*, whose products are known to affect the expression of *stlA* (14) and for genes encoding the global regulators HexA or Hfq.

### Other differentially expressed genes

Other genes affected by IPS treatment encode for products that are involved in competition or defense mechanisms, further suggesting that the bacteria are sensing and responding to stress. In particular, amongst the most highly up-regulated genes are *plu2816* and *plu2817*, involved in the biosynthesis of rhabduscin, a phenoloxidase inhibitor which provides protection against the insect immune system (45). Additionally, genes *plu0884-plu0888, plu1893*-*plu1894, plu4175* and *plu4177* encoding either S-type pyocins or their immunity proteins were up-regulated in the presence of IPS. Furthermore, we observed an up-regulation of genes encoding subunits of the *Photorhabdus* Tc toxins, in particular *tccA1* (*plu4169*), *tccB1* (*plu4168*), *tccC2* (*plu0960*) and *tccC4* (*plu0976*). Finally, we noted the up-regulation of CRISPR related genes such as *plu1791* and *plu1795*, whose products are annotated as a CRISPR-associated nuclease/helicase of the Cas3 type and the CRISPR-associated protein Csy3 respectively.

## DISCUSSION

*Photorhabdus* bacteria have a complex life cycle that includes an insect pathogenic stage and a nematode symbiotic stage. The two stages are associated with distinct metabolic states. More specifically, the virulent stage is associated with active primary metabolism and rapid growth, whilst the symbiotic stage requires secondary metabolism and this shift is reliant on a functional TCA cycle (26). Crucially, mutants in key TCA cycle enzymes are unable to produce stilbenes, light or anthraquinone pigments and are also unable to support IJ recovery in vitro. Importantly, even though these mutants are still virulent towards insects, they give rise to a population within the insect cadaver that has altered characteristics compared to the wild-type bacteria (26). Production of the alarmone (p)ppGpp following nutrient limitation in the insect cadaver, is crucial for survival, induction of infective juvenile recovery and re-establishment of symbiosis (16). *Photorhabdus* commonly exhibits two dominant phenotypic variant types, the primary variant produces light, anthraquinone and other “symbiosis factors” whilst in the secondary variant these factors are absent (46). Both types are virulent but only the primary variant can establish symbiosis. In addition, virulence of the secondary variant was attenuated when secondary metabolism was de-repressed, through mutation of the regulator HexA, even though the ability of the bacteria to form a symbiotic relationship with the nematode was restored (47). It is conceivable that HexA acts to repress premature secondary metabolism, as this would have a detrimental effect on virulence. HexA is a LysR-type regulator with a putative but as yet unknown small molecule ligand (19).

The important role of the TCA cycle in the lifecycle switch of *Photorhabdus* is highlighted by our findings, as the TCA cycle is repressed in the presence of the inappropriate ectopic levels of IPS used. This, and the observation that light and anthraquinone pigment production is inhibited by addition of stilbene to WT *P*. *luminescens* bacteria indicate the presence of a negative feedback loop that controls secondary metabolism. This is also supported by the observation that the *stlA-* mutant appears hyper-pigmented and so likely has deregulated production of anthraquinones (48). The effects of IPS on *Photorhabdus* suggest that the bacteria can actually detect it in a dose dependant manner, as well as produce it. This could either be the result of a direct effect of IPS on the concentration of other metabolic compounds or it could occur through the binding of IPS to one of the LuxR solo receptors. *Photorhabdus* lacks a typical Gram-negative acyl-homoserine lactone quorum sensing (QS) circuit. However it has been shown that one of the LuxR receptors does recognise photopyrones, which are synthesised by *Photorhabdus*, regulating a clumping phenotype (49). Alternatively, IPS could act as the ligand for HexA. However, a previous proteomic study has shown that the TCA cycle is actually up-regulated in secondary variants (50), in which the levels of HexA are higher (19).

In addition to the precise regulation of bacterial metabolism that occurs during the insect infection, it has been demonstrated that following establishment of symbiosis, *P*. *temperata* bacteria residing in the *Heterorhabditis* nematode also experience large transcriptional changes (51). This transcriptional rewiring has the potential of affecting the bacterial metabolism in such a way as to slow down growth and overcome starvation and oxidative damage. This process again needs to be controlled and so far the LysR-type regulator HdfR has been implicated in this (52).

It is of note that the levels of IPS in the *Galleria* infection model remain high for the entire duration of infective juvenile development (21). It has previously been proposed that in addition to its the antimicrobial, antifungal and non-cognate nematode repellent effects, IPS might be acting as the food signal for nematode development. However, it is possible that it is also acting as a signalling molecule for the bacteria to switch to the symbiotic lifestyle. This is supported by the observation that IPS on its own cannot induce IJ recovery (10). It is thus possible that IPS exerts its effects on the bacteria in a way that makes them “primed” for transmission to the nematode and may also induce the release of other compounds that are also necessary for IJ recovery. The actual concentration of IPS may also be important in the effect it has on *Photorhabdus*. Compared to the amounts that can accumulate in the insect cadaver, we used a moderate concentration in this study. We would also like to point out however that it is not known what the intracellular levels of IPS actually are at different stages of the bacterial cycle. This could be modulated by differential changes in membrane permeability or degradation. It is tempting to speculate that mechanisms such as epoxidation or light production might be used as methods to inactivate IPS in the cytoplasm. It was shown for example that an epoxide derivative of IPS has reduced toxicity towards *Photorhabdus* (23). Furthermore, it is known that *trans*-stilbenes, such as resveratrol can easily be converted into the less active *cis* form by exposure to light (53,54).

This would not be the first instance of a compound with antimicrobial properties that can act as a signalling molecule or have an effect on the producer bacteria. It has been actually proposed that the majority of small molecules synthesised by bacteria do indeed play such a role (55). Interestingly, specific acyl-homoserine lactone molecules known to be involved in quorum sensing can also display antimicrobial activities (56). On the other hand, phenazine antibiotics appear to promote survival of Pseudomonas in low oxygen conditions (57) whilst pyocyanin was also shown to be the signal for the activation of genes that are controlled by quorum sensing (58). In summary, we provide evidence of another multipotent small molecule, which is able to modulate the phenotype of the producer bacteria.

## MATERIALS AND METHODS

### Bacterial strains and reagents

The strains used in this study were *P*. *luminescens* subsp. *laumondii* TT01 (WT) (59) a rifampicin resistant derivative of this strain, *P*. *luminescens* spbsp. *laumondii* TT01 (Rif^R^) (60) and a rifampicin resistant *stlA* disruption mutant, BMM901 (TT01 *stlA*::Kn) (24). The strains were grown in Lysogeny broth (LB) or in Lysogeny agar (LBA) plates at 28 °C unless otherwise stated. Purified IPS was a gift from Helge Bode (Goethe Universität Frankfurt, Germany) and additional compound was purchased from iChemical (Shanghai, China). In both cases, purity was confirmed by HPLC.

### Growth, bioluminescence and spectral measurements

Growth (absorbance at 600 nm) and luminescence were monitored using a microplate reader (Omega Fluostar, BMG Labtech). Overnight cultures of *Photorhabdus* were diluted in either Mueller Hinton broth (MH) or LB to an OD_600_ of 0.1. The diluted cultures were then aliquoted in black flat-bottomed µClear^®^ 96-well plates (CELLSTAR®, Greiner) and an equal volume of media containing IPS at double the desired concentration was added to the wells. The plates were then incubated at 28 °C in the microplate reader, with double orbital shaking at 300 rpm and measurements of absorbance at 600 nm as well as luminescence were taken every 30 min. To study the effect of IPS addition during exponential growth, the cultures of *Photorhabdus* were diluted to an OD_600_ of 0.05 and aliquoted in a 96 well plate, 145 µl per well. Growth and luminescence were monitored as described above for 5.5 h until the OD_600_ reached ∼ 0.5. At this point, 5 µl of 960 µg/ml IPS dissolved in DMSO, or DMSO only were added to the wells and the plate was returned to the microplate reader. To compare the levels of luminescence between the two treatments, luminescence was divided by absorbance at 600 nm. Then, multiple t-tests were performed on GraphPad Prism version 8.0.0 (GraphPad Software, available at www.graphpad.com) using the Benjamini, Krieger and Yekuteili method (61) for controlling the false discovery rate. To monitor differences in pigment production, the experiment above was repeated and at the end of the growth curve (t = 24 h) the absorbance of the cultures at the whole spectrum of 250 – 650 nm was measured.

### Biofilm assays

Biofilm was measured using a microplate assay and crystal violet staining. Briefly, bacteria grown overnight were diluted to an OD of 0.05 in LB supplemented with either DMSO or 32 µg/ml IPS. 500 µl aliquots were used to inoculate the wells of a Falcon™ 24-well tissue culture treated polystyrene plate and this was then incubated stationary at 28 °C. After 24 h the supernatant from each well was discarded and the wells were washed three times with PBS to ensure that non biofilm-associated bacteria are removed. The plate was left to dry for 1 h and then 600 µl 0.2 % crystal violet was added to each well. Following a 30 min incubation excess crystal violet was removed by washing the wells three times with PBS. A solution consisting of 70 % ethanol and 10 % isopropanol was used to lyse the cells and release the crystal violet. The amount of dye was quantified using a FLUOstar^R^ Omega microplate reader and measuring absorbance at 570 nm. For each strain, the experiment was performed with four biological replicates and for each of those four technical replicates were used. To assess significance, a paired ratio t-test was performed.

### RNA extraction and sequencing

Overnight cultures of *P*. *luminescens* TT01 were diluted to OD_600_ of 0.05 in two tubes containing 15 ml LB and incubated at 28 °C, 180 rpm until the OD_600_ reached a value of approximately 0.45. Then, to one culture 32 µg/ml of IPS were added, whilst the other culture was treated with an equivalent volume of DMSO. Samples (500 µl) were harvested following incubation for 30 min and treated with RNAprotect (Qiagen). The bacteria were lysed using the enzymatic lysis method described in the RNAprotect handbook in TE buffer containing 20 µl of Qiagen Proteinase K and 1 mg/ml lysozyme. Extraction was performed using the miRNeasy Mini kit (Qiagen) following the manufacturer’s protocol. DNA removal was performed using the Invitrogen TURBO DNA-free kit with Superasin RNAse inhibitor added in the reaction. The samples were then further purified using the Norgen RNA Clean-Up and Concentration kit. The absence of DNA contamination was confirmed by PCR using universal 16S primers 27F (5’-AGAGTTTGATCMTGGCTCAG-3’) and 1429R (5’-CGGTTACCTTGTTACGACTT-3’) (62) and the Agilent Bioanalyser with a Prokaryote Total RNA Pico kit. RNA samples were treated with the bacterial Ribo-Zero rRNA (Bacteria) Removal Kit (Illumina) to remove ribosomal RNA using 2.5 µg of input RNA and 8 µl of Ribo-Zero removal solution per sample. Clean up of depleted RNA was performed using ethanol precipitation. Libraries for sequencing were prepared using the Illumina Truseq Stranded mRNA kit with a slightly modified protocol. Specifically, 5 µl of rRNA depleted RNA were used as input and this was directly mixed with 13 µl of the Fragment, Prime, Finish Mix. Following this, the procedure was as described in the TruSeq Stranded mRNA Sample preparation guide. Sequencing was performed using the Illumina MiSeq Next Generation Sequencer on two MiSeq v3 cartridges with 75 bp paired-end reads. Three sample libraries were sequenced on each cartridge.

### Genome sequencing and Bioinformatic analyses

The genome of *P*. *luminescens* TT01 used in this study was sequenced to account for any differences between lab strains and hence the published genome (GenBank assembly GCA_000196155.1) (63). Sequencing was performed on an Illumina MiSEQ platform with 250 bp paired-end reads. Reads were trimmed using Sickle (https://github.com/najoshi/sickle) and a de novo assembly of the genome was performed with SPAdes v3.0.0 run in ‘careful’ mode (64). Only contigs that were at least 500 bp and had coverage of at least 10 fold were kept for further analysis. The contigs were reordered using Mauve Contig Mover (65) based on the reference assembly GCA_000196155.1. The reordered contigs were artificially joined to create a single fasta sequence that was used as the reference for mapping RNAseq reads. The sequence was annotated using prokka v1.11 (66). Bowtie2 (67) was used for mapping, in ‘very-sensitive’ mode. To identify putative small non-coding RNAs we used the toRNAdo script [28; available at https://github.com/pavsaz/toRNAdo] obtaining a list of ncRNAs for each replicate sample. Putative ncRNAs identified in all three replicates of each condition or in at least 4 samples in total were added to the annotation of the genome. The artificially concatenated and annotated genome is available on figshare (doi: 10.6084/m9.figshare.7473065). Following this, differential expression analysis was performed using DeSeq2 (68) with the file outputs of BEDtools coverageBed (69). A multi factor analysis was performed to measure the effect of IPS treatment whilst accounting for sample pairing. A False Discovery Rate of α = 0.05 was used. To identify the corresponding alleles in the published assembly of *P*. *luminescens* TTO1 (63), a USEARCH global alignment was performed using USEARCH version 8.0 (70) with the protein sequences of assembly GCF_000196155.1 (63) as a database and with a minimum identity threshold of 0.9. Additionally, a whole genome alignment was done using blastn and visualized with the Artemis Comparison Tool (71) to account for any alleles not identified by USEARCH. To investigate the pathways affected, the Uniprot identifiers of proteins encoded by the differentially expressed genes were used as input into STRING database version 10.5 (72) in order to obtain a list of overrepresented KEGG pathways. Multiple testing is corrected for using the method of Benjamini and Hochberg (73). A total of 664 genes could be matched to items in the database. This process was repeated for the up-regulated and down-regulated genes separately, with 422 and 242 items matching to the database respectively.

## AUTHOR STATEMENTS

### Authors and contributions

Alexia Hapeshi: Conceptualisation, Investigation, Formal Analysis, and Writing – Original Draft Preparation.

Jonathan Bennaroch: Investigation.

David Clarke: Resources, Writing – Review and Editing.

Nick Waterfield: Conceptualisation, Project Administration, Writing – Review and Editing, Funding and Resources.

## Acknowledgements

We would like to thank Helge Bode (Goethe Universität Frankfurt, Germany) for providing purified IPS.

