## Supplementary material for "Iso-propyl stilbene: A life-cycle signal?": Figure S1

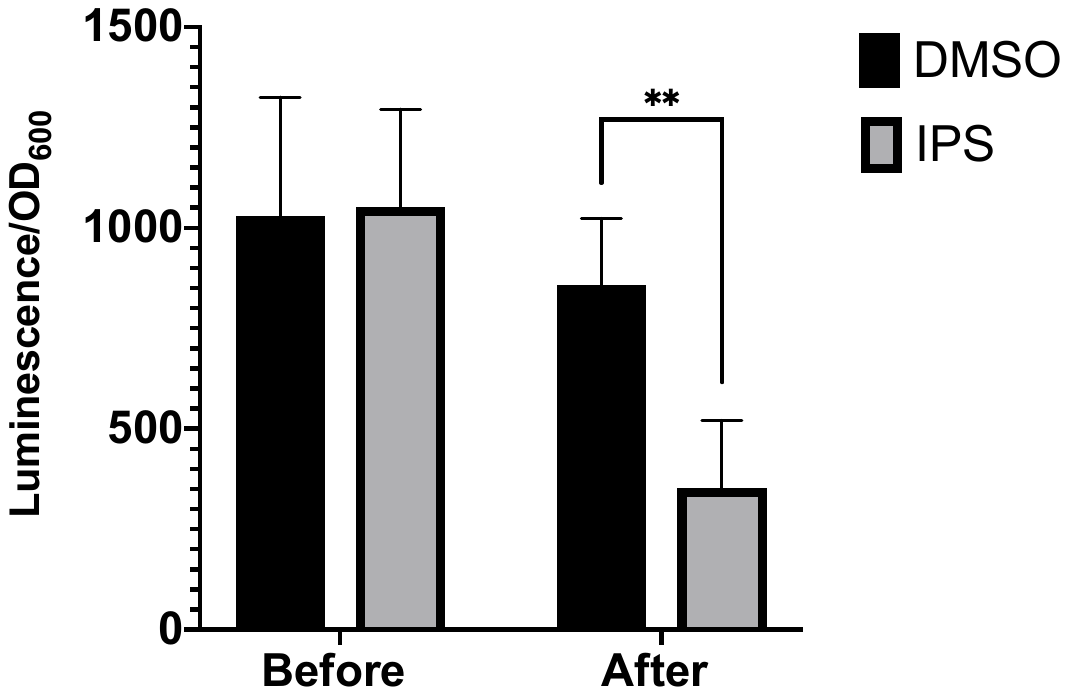

**Figure S1:** Relative luminescence of the cultures sampled for the RNAseq transcriptomic experiment immediately before and approximately 40 min after addition of either 32 µg/ml IPS or DMSO. Results show the averages of the three biological replicates. Asterisks show significant difference at *p* value < 0.01 as measured using a paired t-test.
